# An information theoretic method to resolve millisecond-scale spike timing precision in a comprehensive motor program

**DOI:** 10.1101/2021.07.14.452403

**Authors:** Joy Putney, Tobias Niebur, Rachel Conn, Simon Sponberg

## Abstract

Sensory inputs in nervous systems are often encoded at the millisecond scale in a temporally precise code. There is now a growing appreciation for the prevalence of precise timing encoding in motor systems. Animals from moths to birds control motor outputs using precise spike timing, but we largely do not know at what scale timing matters in these circuits due to the difficulty of recording a complete set of spike-resolved motor signals and relatively few methods for assessing spike timing precision. We introduce a method to estimate spike timing precision in motor circuits using continuous MI estimation at increasing levels of added uniform noise. This method can assess spike timing precision at fine scales for encoding rich motor output variation. We demonstrate the advantages of this approach compared to a previously established discrete information theoretic method of assessing spike timing precision. We use this method to analyze a data set of simultaneous turning (yaw) torque output and EMG recordings from the 10 primary muscles of *Manduca sexta* as tethered moths visually tracked a robotic flower moving with a 1 Hz sinusoidal trajectory. We know that all 10 muscles in this motor program encode the majority of information about yaw torque in spike timings, but we do not know whether individual muscles receive information encoded at different levels of precision. Using the continuous MI method, we demonstrate that the scale of temporal precision in all motor units in this insect flight circuit is at the sub-millisecond or millisecond-scale, with variation in precision scale present between muscle types. This method can be applied broadly to estimate spike timing precision in sensory and motor circuits in both invertebrates and vertebrates.

## Introduction

Neurons in both sensory and motor systems convey information through the precise timing of spikes. Although precise temporal encoding in sensory systems has been found in many organisms [1, 2], the role of temporal encoding in motor systems has been underappreciated. While strong correlations between spike rate and muscle force production exist [3–5], the precise timing of spikes in motor neurons also matters due to nonlinearities in muscle force production and mechanical interactions of musculoskeletal systems within themselves and with the environment [6]. Motor circuits in cortex [7], cerebellum [8], descending interneurons [9], and the periphery [10–12] carry information in neural spike timings. This information can be encoded in the difference between timings in two spike trains [10], the inter-spike interval [7], or temporal patterning [11], providing motor control systems the potential for a rich, layered code with high capacity. Neurons are capable of generating spikes with incredible precision due to biophysical properties that reduce jitter [13], yet this precision may not be consistent with the actual timescale used by the neuromuscular system for motor control. How precisely must the nervous system specify the timing of spikes in the motor periphery to preserve meaningful information about visual stimuli or to control the activation of muscles for movement? Knowing the scale of spike timing precision would inform our understanding of the biophysical properties of muscle and the performance constraints necessary to preserve this precision in the motor periphery.

However, we have few methods to assess at what timescale information about motor output is encoded in motor systems, and how much of the potential bandwidth of temporal codes is utilized. Previous methods for estimating precision include variability on stimulus-response curves [2], spike time jitter analysis in experimental and computational data [14, 15], and information theoretic methods using discrete representations of motor output and response [7]. Using these methods, the precision of firing and encoding of single neurons in some sensory systems has been estimated to occur on the millisecond or sub-millisecond scale [2, 14–16]. In cortical motor circuits, spike timing precision has also been estimated to the millisecond-scale in a songbird vocalization area using a discrete information theoretic method [7]. However, the methods used in these previous studies either do not account for whether precision gives information about an external feature (i.e. a sensory input or motor output) [14] or limit the motor output representation to discrete behavior states instead of utilizing the rich variation that can be present in motor output responses [7]. Therefore, it is difficult to compare spike timing precision in different neural circuits or muscles. Methods that do so would help establish temporal precision in peripheral motor circuits. In motor circuits, the difficulty of obtaining spike resolution is another hurdle to estimating precision. EMG recordings in vertebrates either sample from too many motor units to discriminate spikes or only sample single motor units with spike resolution instead of the entire motor pool, and the calcium dynamics in most imaging techniques occur over greater time-scales than the width of neural spikes.

Hawk moths provide a compelling test-bed for questions of precision due to their fast, complex, and agile motor behaviors that require a rich motor output representation. One hawk moth, *Manduca sexta*, uses a set of only 10 muscles as the primary actuators of their wings and each of these muscles is innervated by one or very few motor neurons, so that each muscle is often considered a single effective motor unit. Therefore, we can record these 10 muscles simultaneously to obtain a nearly complete, or comprehensive, spike-resolved motor program, enabling investigation of precision over a nearly complete circuit that encodes and controls flight. We know that spike timing encodes information about motor output in all muscles of this comprehensive, spike-resolved motor program [12], but do not know the scale of precision utilized for encoding and whether this scale of precision differs in different muscle types. The hawk moth motor circuit, due to its few muscles and spike resolution, enables us not only to estimate timing precision, but discover whether this precision changes depending on muscle type, insertion points, or functional properties across a nearly complete motor program. Being able to estimate precision could point to whether the nervous system encodes movement on a consistent or changing precision scale for different types of muscles.

Here, we take an new approach to estimating spike timing precision in each of the 5 muscle pairs in the hawk moth flight motor program. Our method utilizes the continuous Kraskov *k*-nearest neighbors mutual information (MI) estimation method with additions of uniform noise [12, 17, 18] to corrupt spike timing over progressively larger time windows. We compare this method to a previously established method of estimating spike timing precision using the NSB discrete entropy estimator to evaluate how MI changes with increasingly precise representations of the spiking activity [7, 19]. Our method moves spike timing precision estimates derived from information theoretic methods into the continuous space to give us a more accurate picture of the timescales necessary for encoding motor output. We use this method to assess the scale of spike timing precision in the hawk moth flight motor program, and whether that scale changes in muscles with distinct functional roles.

## Materials and Methods

### Comprehensive, Spike-Resolved Motor Program Data Set

The previously published data set used in this analysis recorded the comprehensive, spike-resolved motor program of the hawk moth, *Manduca sexta* (N = 7), and its motor output in a tethered flight preparation [12, 20]. EMG signals from the 10 primary muscles actuating each moth’s wings were recorded with spike-level resolution using implanted silver wire electrodes (Fig 1A,C). These muscles are treated as single motor units, since each is innervated by one or very few, synchronously firing motor neurons. The moths were tethered using cyanoacrylate glue to a 3D-printed ABS plastic rod attached to a custom six-axis force/torque (F/T) transducer (ATI Nano17Ti, FT20157; calibrated ranges: F*_x_*, F*_y_* = ±1.00 N; F*_z_* = ±1.80 N; *τ_x_*, *τ_y_*, *τ_z_* = ±6250 mN-mm). After tethering, the moths were given thirty minutes to recover from the surgery and adapt to dark conditions, since these are crepuscular moths that typically fly at dusk and dawn. The moths were then presented with a 3D-printed plastic flower oscillating horizontally in a 1 Hz sinusoidal trajectory; these flowers have been used to elicit flight maneuvers previously and sample a wide diversity of turns (Fig 1B) [21]. The EMG recordings were sampled at 10000 Hz, amplified using a 16 channel amplifier (AM Systems Inc., Model 3500), and acquired using a data acquisition board (National Instruments USB-6529 DAQ) and custom MATLAB software. The same DAQ board was used to acquire the strain gauge voltages from the ATI F/T transducer used to calculate the forces and torques, also sampling at 10000 Hz.

**Fig 1.**
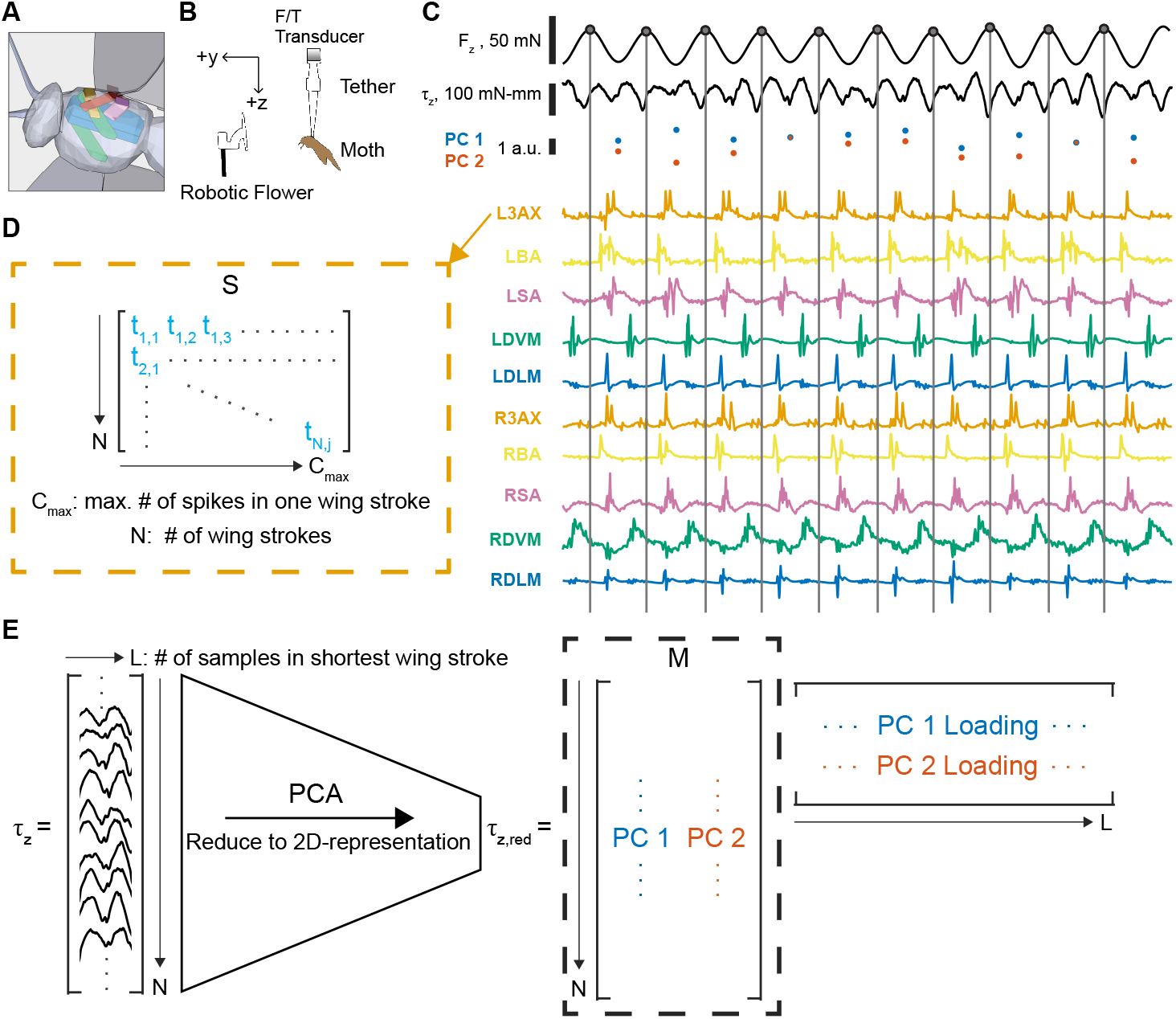
Representing the spiking activity and motor output of a moth’s comprehensive, spike-resolved motor program. A: A 3D schematic showing approximate positions and attachments of the 5 muscle types recorded in the motor program: the dorsolongitudinal muscle (DLM, blue), the dorsoventral muscle (DVM, green), the third axillary muscle (3AX, orange), the basalar muscle (BA, yellow), and the subalar muscle (SA, pink). B: Tethered preparation used to record EMG signals from the 10 muscles in the motor program and the forces and torques produced as the moth responded to a flower stimulus. C: Example data from 0.5 seconds of tethered flight in a moth. The bandpass-filtered *F_z_* signal is used to segment wing strokes, with t = 0 of every wing stroke corresponding to the peak downward force in *F_z_* (gray circle and line). Spike timings in each muscle and the yaw torque *τ_z_* are aligned to t = 0 within each wing stroke. D: The spiking activity *S* for each muscle and each moth can be represented as a *N* × *C_max_* matrix where *N* is the number of wing strokes sampled in that moth and *C_max_* is the maximum spike count observed in that muscle. Where there are fewer than *C_max_* spikes in a wing stroke, entries with no spikes are represented as *NaN*. E: The yaw torque *τ_z_* after wing stroke segmentation can be represented as a *N* × *L* matrix where *N* is the number of wing strokes sampled in that moth and *L* is the length in samples of the shortest wing stroke in the data set, meaning that some wing strokes are shortened to length *L*. The matrix *τ_z_* can be reduced to a *N* × 2 matrix using principal components analysis (PCA). The motor output *M* is represented as the projection of each wing stroke onto the first two principal component (PC) loadings that explain the variance of the fully dimensional *τ_z_*. Data shown and panels A-B taken and adapted from Putney et al. 2019 [12, 20].

The data set reports spike counts and spike timings in segmented wing strokes as representations of the spiking activity along with the scores of the first 2 principal components (PCs) of the within-wing stroke yaw torque (*τ_z_*) produced in those wing strokes each individual moth (Fig 1D-E). The wing strokes were segmented using a previously described method [22]. The force in the z-axis (F*_z_*) was filtered with a 8-th order Butterworth bandpass filter between 5 and 35 Hz, capturing the wing beat frequency of the moth. A Hilbert transform was used to identify the peak downward F*_z_* in each wing stroke to serve as the zero time point, t = 0, for each wing stroke. The timing of spikes in each muscle within each wing stroke were aligned to these zero time points, such that the spiking activity were represented as continuous times in ms within each wing stroke as a matrix *S* (Fig 1D, S1 Fig). The within-wing stroke yaw torque was also segmented into wing strokes using the identified zero time points, and a principal components analysis of the continuous yaw torque signal from t = 0 to the maximum length *L* in samples of the shortest wing stroke of each individual moth was conducted (Fig 1E). The scores – the projections of each wing stroke onto the first two principal components – were used as a motor output representation *M* of the within-wing stroke yaw torque, *τ_z_*. For full details on the wing stroke segmentation and PCA of the yaw torque, see the original paper [12].

To estimate spike timing precision, we first adapted a previously published discrete information theoretic method that utilizes the Nemenman-Shafee-Bialek (NSB) entropy estimator to determine the mutual information (MI) between discrete probability distributions of spike “words” and motor output states [7, 19, 23, 24] (Fig 2). Here, spikes are represented discretely by binning the spikes within the wing stroke into smaller and smaller bins, providing a more precise representation of where spikes occur within the wing stroke. We then introduce a new approach that builds off the Kraskov-Stögbauer-Grassberger (KSG) continuous mutual information estimator used to determine the MI between spike timings and motor output to estimate spike timing precision by adding uniform noise in varying window sizes to the spike timings [12, 17, 18] (Fig 3). This method is only limited in its representation of the spike timings by the sampling rate. With a sampling rate of 10000 Hz, spike timings are represented to 0.1 ms.

**Fig 2.**
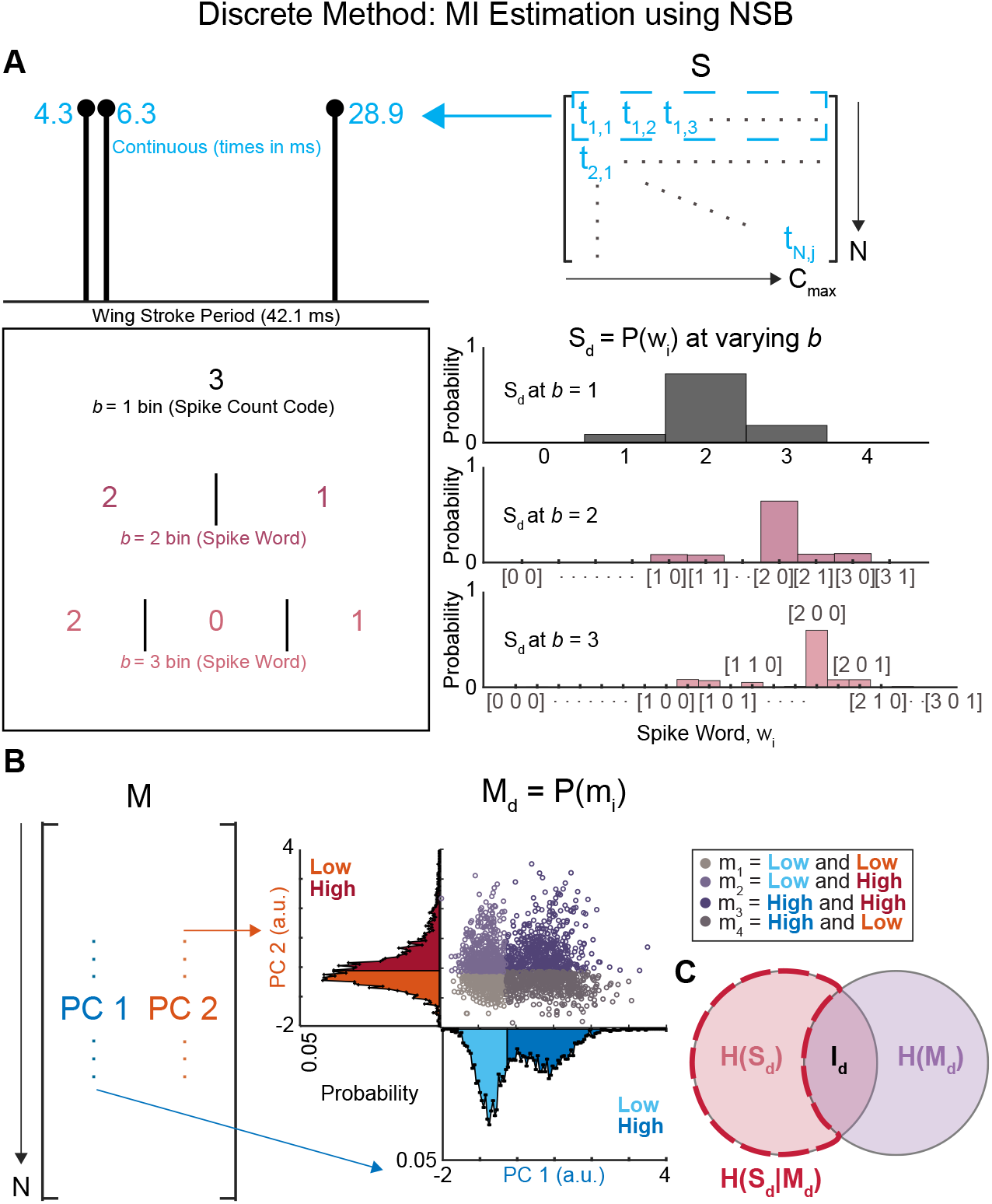
Discrete Method: MI Estimation using NSB. A: The spike train within a wing stroke can be represented by a pure continuous code with the precise spike timings (blue), a pure spike count code where the activity is represented just by the number of spikes in the train (black), or binned codes that contain more precise information with more bins (gray). The spiking activity representation *S* is transformed into a discrete probability distribution *S_d_* whose entropy can be estimated using the NSB method. Each spike train in a wing stroke is binned into spike words *w_i_* with *b* bins. The probability distribution of occurrences of each spike word *w_i_* is our representation of the spiking activity as a discrete probability distribution. B: The motor output represention *M* is transformed into a discrete probability distribution *M_d_*. The scores of the first two principal components can be used to specify four behavioral states *m_i_* where *i* = 1-4 categorized by low and high groups above and below the median score for each component. Visualization of the four motor output states *m_i_* as a scatter plot of the scores of the first two principal components, with each unique color specifying a motor output state. C: The NSB entropy estimator is used to estimate the entropies of the spiking activity *H*(*S_d_*) and the conditional entropies *H*(*S_d_*|*M_d_*) at different numbers of bins *b* to determine the mutual information (*I_d_*) between the spiking activity *S_d_* and the motor output *M_d_*.

**Fig 3.**
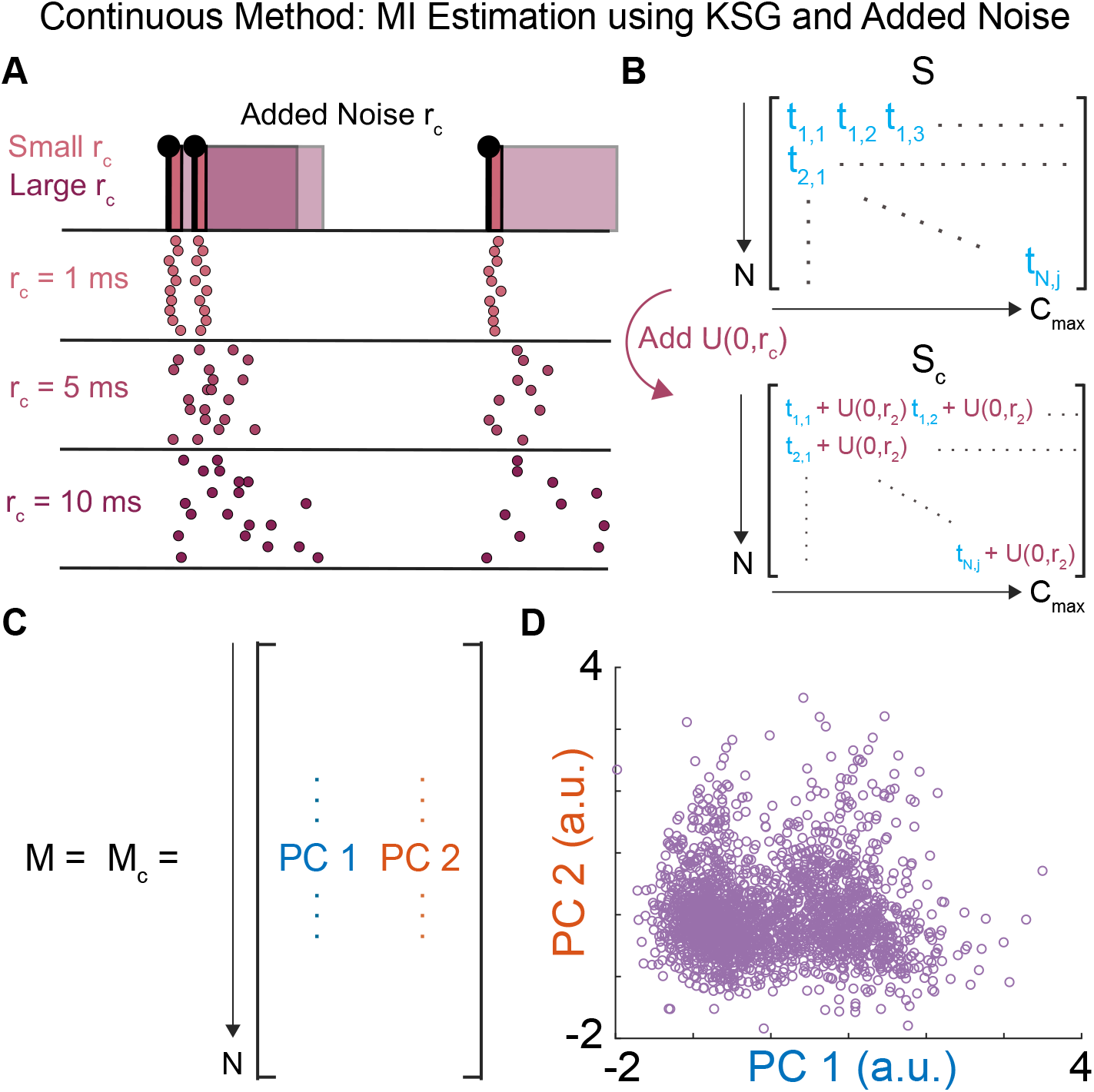
Continuous Method: MI Estimation using KSG and Added Noise. A: Uniform noise (*U*(0, *r_c_*) is added to the precise spike timings. At small *r_c_*, the noise corruption does not change the spike train representation much over 10 iterations. At large *r_c_*, significant variation between the representation of the same spike train appears over 10 iterations. B: The representation of the spiking activity *S* has a uniform window of noise defined by width *r_c_* added to it to create representations *S_c_* at varying levels of added noise. C: The motor representation for the continuous method *M_c_* is the same as the continuous representation described above *M* which is a *N* × 2 matrix of the scores of the first two principal components for each wing stroke. D: Visualization of *M* = *M_c_* as a scatter plot of the scores of the first two principal components, where each point is a wing stroke.

### Discrete Method: MI Estimation using NSB

We first attempted to estimate spike timing precision using a previously described NSB entropy estimation method [7, 19, 23, 24].The spiking activity was used to create probability distributions *S_d_* with discrete states of spike “words” *w_i_* created by binning the spikes in each wing stroke using different numbers of bins, *b* (Fig 2A). The number of spikes that occur in each bin sets the value of that bin, and each unique sequence of bin values across all wing strokes is a spike “word” *w_i_*. The prevalence of each spike word provides a discrete probability distribution *S_d_* = *P*(*w_i_*) that describes the spiking activity in each individual muscle. The number of bins *b* used to create this discrete PDF was varied; the size of each of these bins set the spike timing precision *r_d_* as:

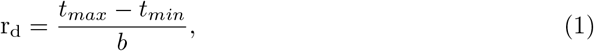

where *t_max_* is the highest spike time recorded in that muscle and *t_min_* is the lowest. This is the maximum range within a wing stroke where spikes occur. When this range is split into *b* bins the size of each bin is *r_d_* in milliseconds.

To create a discrete probability distribution for the motor output, we used the median of the PC scores *M* on each axis (each column of *M*) to create behavioral groups delineated by low and high states where values below the median were placed into the “low” behavioral group and values above the median were placed into the “high” behavioral group (Fig 2B). This allowed us to then create a discrete probability distribution *P*(*m_i_*) of 4 motor output states where wing strokes were categorized as being “low” or “high” on both axes, or “low” on one axis and “high” on another axis. The motor output representation *M_d_* then is the discrete probability distribution *P*(*m_i_*) where *m_i_* are each of four motor output states (*i* = 1-4). In a previous implementation of this method to songbird motor cortex data, the motor output was divided into only 2 potential states using the method above [7]. Here, we use 4 motor output states to allow us to draw comparisons with the continuous method described below which uses scores from both principal components and to maintain consistency with the previously published paper on this data set which used the scores from both principal components to estimate the mutual information between spiking activity and motor output [12].

Using these two discrete probability distributions, we estimated the mutual information (MI) between the spike “words” (represented by *S_d_*) and the motor output states (represented by *M_d_*) for each muscle in each moth (Fig 2C):

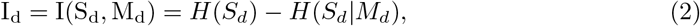

where the mutual information *I_d_* = *I*(*S_d_*,M*_d_*) between the spike word probability distribution *S_d_* and the motor output state probability distribution *M_d_* is equal to the entropy of the spike word probability distribution (*H*(*S_d_*) minus the conditional entropy of the spike word probability distribution given knowledge of the motor output state *H*(*S_d_*—*M_d_*). This conditional entropy is the sum of the entropy of the spiking activity for wing strokes in each motor state, *m_i_*, weighted by the probability of that motor state *p*(*m_i_*):

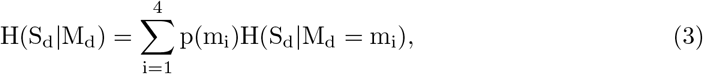

Since all entropy estimates with finite data have bias, we used the NSB entropy estimation for the entropy of the full spike word distribution *H*(*S_d_*) and the conditional entropy *H*(*S_d_*|*M_d_*) in this equation [7, 19, 23, 24], which can estimate these entropies with less bias even when severely undersampled as opposed to the more traditional direct method, or maximum likelihood, estimation [1]. The NSB method uses Bayesian estimation of the underlying probability distributions given observed events, and then estimates the first and second posterior moments of the entropies. The reported mean and standard deviation of these entropy estimates was calculated using the first and second posterior moment of the entropy, respectively [24]. Estimates of *I*(*S_d_*, *M_d_*) were done for spike word distributions *S_d_* defined with increasing numbers of bins *b* = 1 - 70, with *b* = 70 being well within the sub-millisecond range in all muscles. With this method, if a plateau in MI can be identified as *b* increases, a breakpoint analysis could be used to determine the spike timing precision *r_d_* by fitting a two-region regression to the plateau and the region where MI is increasing with increasing *b*; the breakpoint between these two regions would be the spike timing precision *r_d_*.

### Continuous Method: MI Estimation using KSG and Added Noise

Our second method of estimating the spike timing precision used a continuous Kraskov *k*-nearest neighbors MI estimation [17] (Fig 3). The Kraskov *k*-nearest neighbors method uses the Euclidean distances between each sampled wing stroke and its *k*th nearest neighbor in a space defined by variables *X* and *Y* to estimate the joint entropy of the two variables *H*(*X*, *Y*) and the mutual information between the two variables *I*(*X*, *Y*). *X* and *Y* can be multidimensional, with size *N* × *v* where *N* is the number of observations and *v* is the number of dimensions that define the variable *X* or *Y*. This method also preserves continuous information since no discretization of the variables *X* or *Y* is required. Here, *X* is a representation of our spike timings, *S_c_*, which has dimension size *N* × *C_max_* (Fig 3B). *Y* is the scores of the two principal components of the yaw torque (*M* = *M_c_*), which is a 2-dimensional continuous variable (Fig 3C). For the spike timings, we generated a random uniform distribution of noise *U* between values of 0 and *r_c_*, where *r_c_* is the upper limit of the distribution (Fig 3A). This adds a window of noise equivalent to a value *r_c_*, in milliseconds, shifting spike timing values *S_c_* by *U*(0, *r_c_*) (Fig 3B). We then estimate the MI between the noise corrupted spike timing values and *M_c_* across values of *r_c_*:

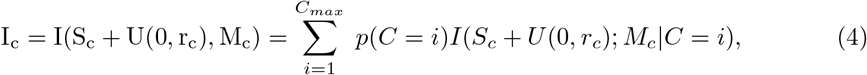

where *C* is the number of spikes in a wing stroke and *C_max_* is the maximum count of spikes in a single wing stroke for each muscle. This formulation for the information between spike timings and the motor output allows for a variation in the number of spikes in each wing stroke to be correctly taken into account. The mutual information between the spike timings with added noise *S_c_* and the scores of the first 2 PCs in each wing stroke *M_c_* was estimated 150 times with different random noise samples for each value of *r_c_* tested, reducing the effect any individual estimation run has on the final mutual information value. Therefore, we report the mean and standard deviation of these 150 estimates of *I_c_*.

We defined the precision as the point where the MI estimate falls below the lower bound on the estimate of the MI (the mean minus the standard deviation) at *r_c_* = 0.00 ms (i.e. the original data) based on variance in data fractions [12, 18]. The code implementation of the KSG estimator we used does jitter the values of the variables, but at several orders of magnitude below the level of uniform noise we add. We determined the standard deviation of our MI estimates at *r_c_* by subsampling the data sets in non-overlapping data fractions. The variance in these fractions was used to estimate the standard deviation or uncertainty of the estimate at the full data size, as previously [18]. To test for statistically significant differences between the spike timing precision *r_c_* of different muscles, we used one-way ANOVA and Kruskal-Wallis tests.

## Results and Discussion

### The continuous method gives an improved estimate of millisecond-scale timing precision

Here, we present an extension of a continuous MI estimation method [18] to obtain reliable estimates of millisecond-scale timing precision (Fig. 4, 5) across the hawk moth’s comprehensive motor program. The continuous method with the disrupting role of additive noise allows us to determine a specific estimate of the spike timing precision *r_c_* at fine resolution, giving an improved assessment of spike timing precision over the discrete method. The value of added noise *r_c_* where the mean MI estimate dropped below the range of error for *I_c_* at *r_c_* = 0 was chosen as the threshold for spike timing precision.

**Fig 4.**
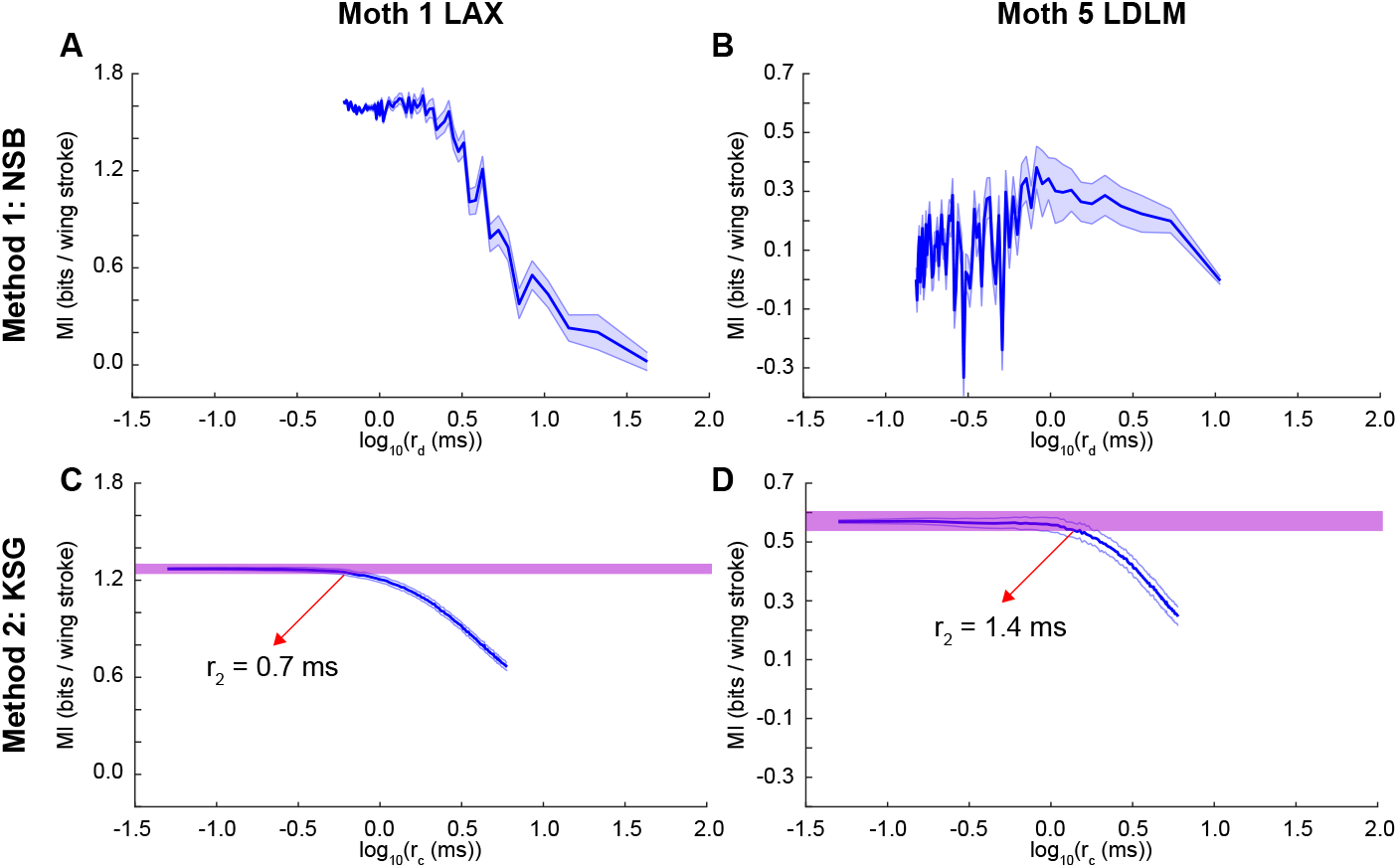
Comparing the two methods used to estimate spike timing precision. A-B: MI (mean ± STD) estimated using the Discrete Method for a sample L3AX muscle (A) and a sample LDLM muscle (B). C-D: MI (mean ± STD of 50 tests of adding uniform noise) estimated using the Continuous Method at values of uniform noise added for the same L3AX muscle as in A (C) and the same LDLM muscle as in B (D) (red arrow: estimate of spike timing precision, *r_c_*; purple line: error estimate of MI at zero noise added). All values are plotted on a log scale.

**Fig 5.**
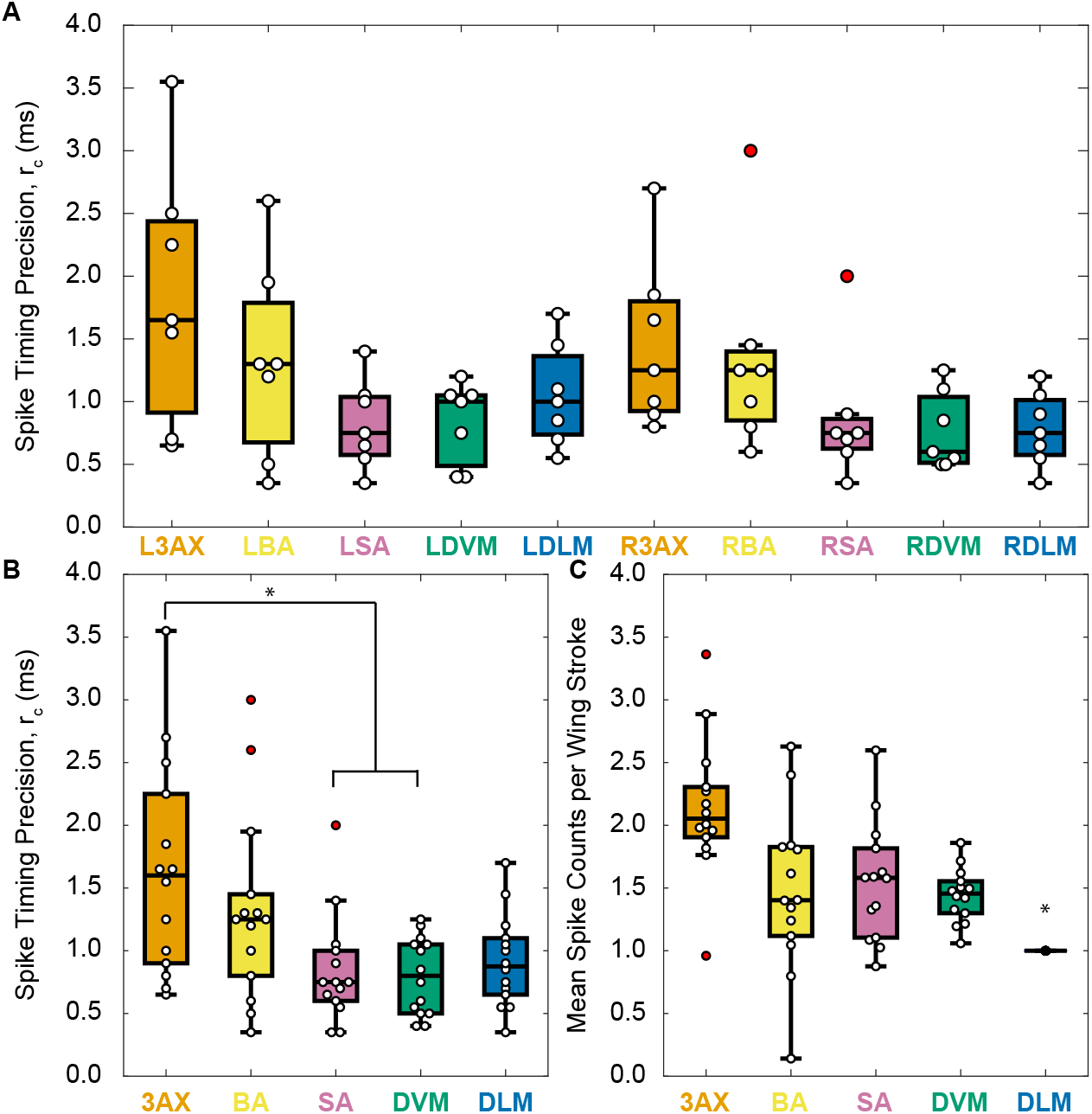
Spike timing precision estimates from the Continuous Method. A: Spike timing precision values (*r_c_*) for all moths reported as boxplots (middle black line: median; boxes: 25th and 75th percentiles; whiskers: all data except outliers; white circles: *r_c_* values for individual moths; red circles: *r_c_* values for moths that were considered statistical outliers). Marginally statistically significant differences were found across muscle types (one-way ANOVA, p = 0.0147; Kruskal-Wallis, p = 0.0655). B: Spike timing precision values (*r_c_*) for all moths with left and right side muscles combined (same data as A). The asterisk denotes groups that with statistically different means in a Kruskal-Wallis non-parametric test. The precision values of the 3AX muscle is significantly different from the SA and DVM muscles in a multi-comparison Kruskal-Wallis test at significance level p ¡ 0.05 with corrections using the Tukey’s honestly significant difference criterion (p = 0.0057). The precision of the 3AX muscle was significantly different from the SA and DVM muscles in a multi-comparison two-way ANOVA with the same significance criterion as for the Kruskal-Wallis test (two-way ANOVA for muscle type, p = 0.0012; and left/right sides of the moth, p = 0.36; with a test for interaction, p = 0.85). C: Mean spike counts for all moths with left and right muscles combined as in B. The asterisk denotes groups with statistically different means in a Kruskal-Wallis non-parametric test. The spike count means of the DLM was significantly different from all other muscles in a multi-comparison Kruskal-Wallis test at the same significance criteria as in B (p < 10^−6^). The 3AX muscle is significantly different from all other muscles and the DLM is significantly different from the SA muscle in a multi-comparison two-way ANOVA with the same significance criterion (two-way ANOVA for muscle type, p < 10^−6^; and left/right sides of the moth, p = 0.62; with a test for interaction, p = 0.93).

The discrete method, which has been previously used to show spike timing precision in motor systems [7, 24], could not provide a consistent precision estimate in all muscles. One of the advantages of the NSB estimator over other discrete estimators is that it can estimate probability distributions from very sparse sampling. However, in our data set, this method fails to establish a lower bound on precision. Some estimates were unstable at high precision, as seen in Moth 5’s LDLM (Fig 4B), where negative information values were obtained at high numbers of bins. Others, even when sampling over 70 bins in a wing stroke, did not reach the precision obtainable by the continuous estimation method, as seen in Moth 1’s L3AX (Fig 4A). Still other examples never plateau at high bin sizes (S2 Fig). In contrast, the continuous noise addition method resulted in stable estimates, with an expected monotonic decrease in MI as the amount of added uniform noise increased that was consistent across all muscles in our data set (Fig 2C-D, S3 Fig). Using the continuous method, we can estimate the mutual information at smaller values of precision and have greater stability at these smaller values (Fig. 4).

Our implementation of the discrete method estimated mutual information for a motor output probability distribution *M_d_* that included more motor output states *m_i_* than its previous implementation, which only predicted between two motor output states [7]. Simplifying *M_d_* to only include two motor output states did not significantly improve the quality of our estimates. In other motor circuits like those found in cortex, it might be possible possible to obtain enough spike trains to produce stable estimates by more completely sampling the range of all possible realizations of spiking activity, since this method produced stable estimates when implemented previously [7]. Even still, the continuous method still provides several advantages over the discrete method. With the continuous method, we do not have to discretize the motor output and therefore can estimate MI with a much richer representation of the motor output space. For example, in Moth 1’s L3AX muscle (Fig 4A,C), a plateau in MI occurs at a larger precision value in the discrete method than the continuous method; however, the motor output representation in the discrete method is much simpler, consisting of four motor output states. With the continuous method, we incorporate a continuous representation of the motor output and see a much smaller precision value, highlighting the importance of within-wing stroke features of the motor output that are not captured in the four motor output state space in the discrete method. Additionally, in our spiking activity representation, we also have the advantage of being able to more finely represent different levels of spike timing precision, especially at small values of *r_c_*, whereas the discrete method is limited to binning spike trains. This enables us to determine a specific estimate of spike timing precision at fine scales with the continuous method. Finally, the KSG MI estimation only includes information encoded by spike timings, whereas the NSB MI estimation includes information about spike rate in its entropies; this feature of KSG has been used to separate spike rate and timing information [11, 12].

The discrete method has been used to show millisecond-scale encoding in a cortical vocal area in songbirds [7]. While this method is not fully stable or rich enough to confidently resolve the degree of precision it is generally consistent in showing millisecond-scale precision in hawk moth flight muscles, in that estimates of mutual information is increasing in most cases where a smooth trend is still observed (S2 Fig). Our implementation of noise addition allows for more confident estimates of spike timing precision using a continuous information theoretic method in experimental data, especially at sub-millisecond scales. Other methods besides those presented here have been used to estimate spike timing precision. Many experimental studies in sensory systems have investigated the statistics of spike timing jitter as a proxy for how precise the neural system specifies must be to encode information about a repeated stimulus [25–27]. In motor systems, however, we cannot obtain many realizations of the same motor output to investigate jitter in spike timings because it is difficult to constrain motor behavior. Even in well-defined tasks like reaching along a pre-defined trajectory, variation exists in how a motor task is realized [28, 29]. Computational studies have also used spike timing jitter to measure the reliability of neural coding under repeated presentations of the same sensory stimulus [14, 30]. For motor systems, however, other methods are necessary because the moth will not produce the same motor output repeatedly to estimate spike timing jitter. Jittering methods can also fail to address whether the neuronal precision observed is actually necessary for encoding either the sensory stimulus or motor output. While we have known that computational modesl of noisy neurons in Kilinc et al. can demonstrate precisely time spikes [30] and neural networks can learn precisely timed sequences of spikes [31], we can now assess the degree of spike precision across the entire comprehensive motor program. The continuous method developed here can be used for both sensory and motor systems to determine the scale of spike timing precision at fine resolution.

One implementation of spike timing jitter used Gaussian noise windows in data from LGN neurons to estimate mutual information between spike trains and visual stimulus movies, which does address whether spike timing precision informs motor output [15]. Here, instead of Gaussian noise, our method uses uniform noise to evenly jitter the spike timings over a time window that can correspond more directly to the bin sizes used in the previously published discrete method. Additionally, our method of noise addition preserves no information about when the spike occurred, whereas adding a Gaussian window of noise will still preserve that information because the distribution of jittered spike times will peak at the original spike timing. Using a uniform window of noise corrupts information evenly across the window *r_c_* which we use to define the spike timing precision.

### Every muscle in the hawk moth flight motor system uses millisecond-scale spike timing precision

Using the continuous method, we demonstrate spike timing precision to the millisecond-scale across all ten muscles in the motor system as well as statistically significant differences in the spike timing precision of functionally distinct muscle types (Fig 5A). When left and right muscles are considered separately, there are marginally statistically significant differences across the muscle groups, but all mean values across individuals and muscles range from 0.8 to 1.8 ms. We combine left and right muscles of each type to determine whether different muscles have statistically different spike timing precision values (Fig 5B).

Each of the five muscle types in the hawk moth motor program – the dorsolongitudinal (DLM), dorsoventral (DVM), third axillary (3AX), basalar (BA), and subalar (SA) muscles (Fig. 1A,C) – have different sizes, different attachment points, different activation mechanisms, and different functional roles. The DLM and DVM indirectly actuate the down stroke and up stroke of the wing, respectively, by deforming the thorax. The 3AX, BA, and SA muscles directly attach to the base of the wing to fine-tune the motion of the wing during flight. The DLM and DVM have been traditionally thought of as flight power muscles, while the 3AX, BA, and SA have been called flight steering muscles [12]. In all of muscles, the importance of spike timing for motor output has been supported. Sub-millisecond scale timing in the DLM is linked to power output during flight [10, 32]. Changes in the spike timing of the DVM and 3AX in a sister species of hawk moth to the one we investigate here have been shown to correlate with wing kinematic changes at wing stroke reversal [33]. In *Manduca sexta*, changes in timing of the BA is correlated with turning behavior and changes in the timing of the SA is correlated with wing depression, promotion, and remotion [34, 35]. Because each of these muscles is functionally distinct, we investigated whether the scale of spike timing precision differed by muscle type.

The lowest mean spike timing precision of a muscle (*r_c_*) across all moths is observed in the subalar (SA) and dorsoventral (DVM) muscles, with mean precision values of 0.8 ms, which makes them on average sub-millisecond precise. We also found the DLM muscles to be sub-millisecond precise on average. Previously, bilateral timing differences between the left and right DLMs were shown causally to have sub-millisecond precision for producing yaw torque [10]. We can now demonstrate that both sets of flight power muscles, the DLMs and DVMs, encode information on the sub-millisecond scale as individual muscles, not just in relation to each other. The ability to fine-tune power production during flight to produce yaw turns is driven by sub-millisecond changes in these power muscles, and sub-millisecond differences in their timings relative to each other. This rebuts the traditional view that these flight power muscles do not have fine-tuned control of flight. Additionally, evidence of sub-millisecond scale encoding in the SA muscles points to a precise control role for these muscles, perhaps in controlling wing depression and promotion/remotion at its specific phase of firing [34].

The 3rd axillary (3AX) muscles have the largest mean spike timing precision at 1.6 ms and are statistically different from the SA and DVM muscles in a multi-comparison Kruskal-Wallis test. This demonstrates the absence of a link between the level of spike timing precision and amount of spike timing MI encoded by each muscle type, since the muscles have been previously shown to encode about the same amount of spike timing information [12]. While all these muscles encode the same amount of spike timing information, they have different overall amounts of variation in spike timing during a wing stroke (S1 Fig). The 3AX muscle is able to encode the same amount of information about the motor output, but does so with less spike timing precision. While not statistically significant, the mean spike count of the 3AX muscle is higher than all other muscle types in the motor program (Fig 5C). It could be that having multiple spikes to carry motor information decreases precision requirements for each spike. However, the DVM and SA both report lower mean precision values than the DLM despite having more spikes. The DLM remains one of the most precise muscles in the motor program though, which may be driven by the constraint of having only one spike with which to encode information.

The spike timing precision of individual muscles may also enable coordination between muscles. The studies cited above that demonstrate the importance of timing throughout the hawk moth motor program all reported spike timings of muscles relative to the DLM [32–35], and the study on the DLM timing investigated the relative timing between the left and right DLMs [10]. Here, we demonstrate that all individual muscles have precision on the sub-millisecond to millisecond-scale, which could provide the precision necessary for this type of coordination across muscles.

### Spike timing must be essential for sensorimotor integration

Because millisecond-level spike timing precision is present across the entire motor program, not just in specialized muscles, encoding of this precision must either be preserved throughout the nervous system from sensory inputs to interneurons then out to the periphery, or could be accomplished entirely in the periphery via direct sensorimotor connections. Many sensory systems are millisecond-scale precise [2], much like the flight motor program investigated here, so sensory inputs are likely candidates for encoding information precisely that could then be preserved in the transformations of activity to muscles. In the hawk moth investigated here, *M. sexta*, it is known that dopaminergic interneurons whose activity is correlated with visual stimulus changes heavily innervate the pterothoracic ganglion with axon branches where the motor neurons controlling the 5 muscles studied here originate [36, 37]. These interneurons could preserve temporally precise information about the visual scene that is passed through synapses to produce precise spiking. Dopaminergic interneurons that synapse directly with the motor neuron for the basalar muscle (b1) in flies enable wing coordination during onset and termination of flight by integrating bilateral sensory inputs [38]. Additionally, precise timing in a GF interneuron relative to parallel circuits has also been found to determine action selection in a Drosophila escape response [9]. This demonstrates the importance of precise timing in the central nervous system for action selection – a direct effect on motor behavior – and may point to the preservartion of precision throught a sensorimotor circuit since these GF interneurons receive input from the optic lobes.

An alternative possibility is that precision arises not from precise visual encoding preserved through descending interneurons, but from direct sensorimotor connections in the periphery from mechanosensors. In flies, activation of haltere steering motor neurons is known to modulate the spike timing of a basalar muscle on the millisecond-scale [39], facilitated by a direct, monosynaptic connection between haltere afferents and the motor neuron of that muscle [40]. Haltere mechanosensors are typically campaniform sensilla, which have been shown to encode lateral displacements using precise spike timing [41]. While moths do not have halteres, they do have campaniform sensilla on their wings that respond within milliseconds to mechanical stimulation [42], which could dictate temporally precise responses in muscles. Flight posture reflexes have been activated by stimulating these wing mechanosensors [43], which may indicate the presence of analogous direct monosynaptic connections between wing mechanosensors and motor neurons in the hawk moth. It is likely that both these peripheral sensorimotor connections as well as descending inputs carrying visual information play a role in preserving spike timing precision in the motor periphery that is necessary for the robust, agile execution of flight.

While previously sub-millisecond precision had only been demonstrated in a bilateral pair of muscles in the hawk moth flight motor program, we now have shown at least millisecond-level encoding precision in each individual muscle that the moth uses to control its wings during flight. The presence of highly temporally precise coding is not a special case reserved for certain functionalities, but rather a pervasive encoding strategy used by the nervous system. Precision is likely necessary due to large changes in power output driven by millisecond-scale changes, which belies the assumption that muscle can be treated merely as a low pass filter on motor neuron activity, where spike rate is proportional to force produced. Millisecond changes in spike triplets in motor units of song bird breathing muscles causally change pressure production in their lungs [11]. The biomechanical and molecular properties of muscle change throughout their cycle of activation, and the millisecond scale timing of activation during these state changes could cause muscle to produce different forces. While hawk moth flight is a relatively “fast” behavior, with wing strokes elapsing 40 to 50 milliseconds, temporal encoding should not be assumed to only occur in insects or other animals with fast frequency behaviors. Walking or running can be a much lower frequency behavior, but bipedal foot strikes – which occurs on a similar timescale to a moth wingstroke – may use precisely timed motor signals [44]. Slower time scale behaviors, like breathing in songbirds, have also been shown to encode on the millisecond scale [11]. Other motor behaviors in vertebrates also can occur on fast timescales, like eye movements, finger snapping, and typing. Additionally, neural mechanisms in vertebrates could be used to encode and preserve spike timing precision across synapses. As an example, stretch reflexes use direct sensory-to-motor connections that could be analogous to the highly specialized circuits in insect flight, and may serve as sources of precision in vertebrate motor units. The spike timing precision of motor neuron activity in a variety of behaviors and species can be assessed using the method presented here, which provides a specific estimate of the scale of precision for encoding a motor output.

## Supporting information

**S1 Fig. Raster plots from five muscles along with corresponding PC scores.** Timing of the L3AX, LBA, LSA, LDVM, and LDLM muscle spikes for 200 example wing strokes in an individual moth (scale bar = 20.0 ms). Spikes are color-coded by their order in each wing stroke. The corresponding values in arbitrary units of the PC scores in each wing stroke are displayed in the far right panel.

**S2 Fig. Examples of estimates of** *I_d_* **from the Discrete Method using NSB with four motor states** *m_i_* **where** *i* = 1 – 4. Log plots of *I_d_* at different values of *r_d_* for three moths and three muscle types. The axes here should correspond to the ones in S3 Fig for comparison of the methods.

**S3 Fig. Examples of estimates of** *I_c_* **from the Continuous Method using KSG.** Log plots of *I_c_* at different values of *r_c_* for three moths and three muscle types. The axes here should correspond to the ones in S2 Fig for comparison of the methods.

## Acknowledgments

This material is based upon work supported by a NSF Graduate Research Fellowship (DGE-1650044) awarded to J.P., an NSF Faculty Early Career Development Award (Award no. 1554790) to S.S., and a Klingenstein-Simons Fellowship in the Neurosciences to S.S.

