## Supplementary figures and images for "An information theoretic method to resolve millisecond-scale spike timing precision in a comprehensive motor program"

### Supplemental Figure 1

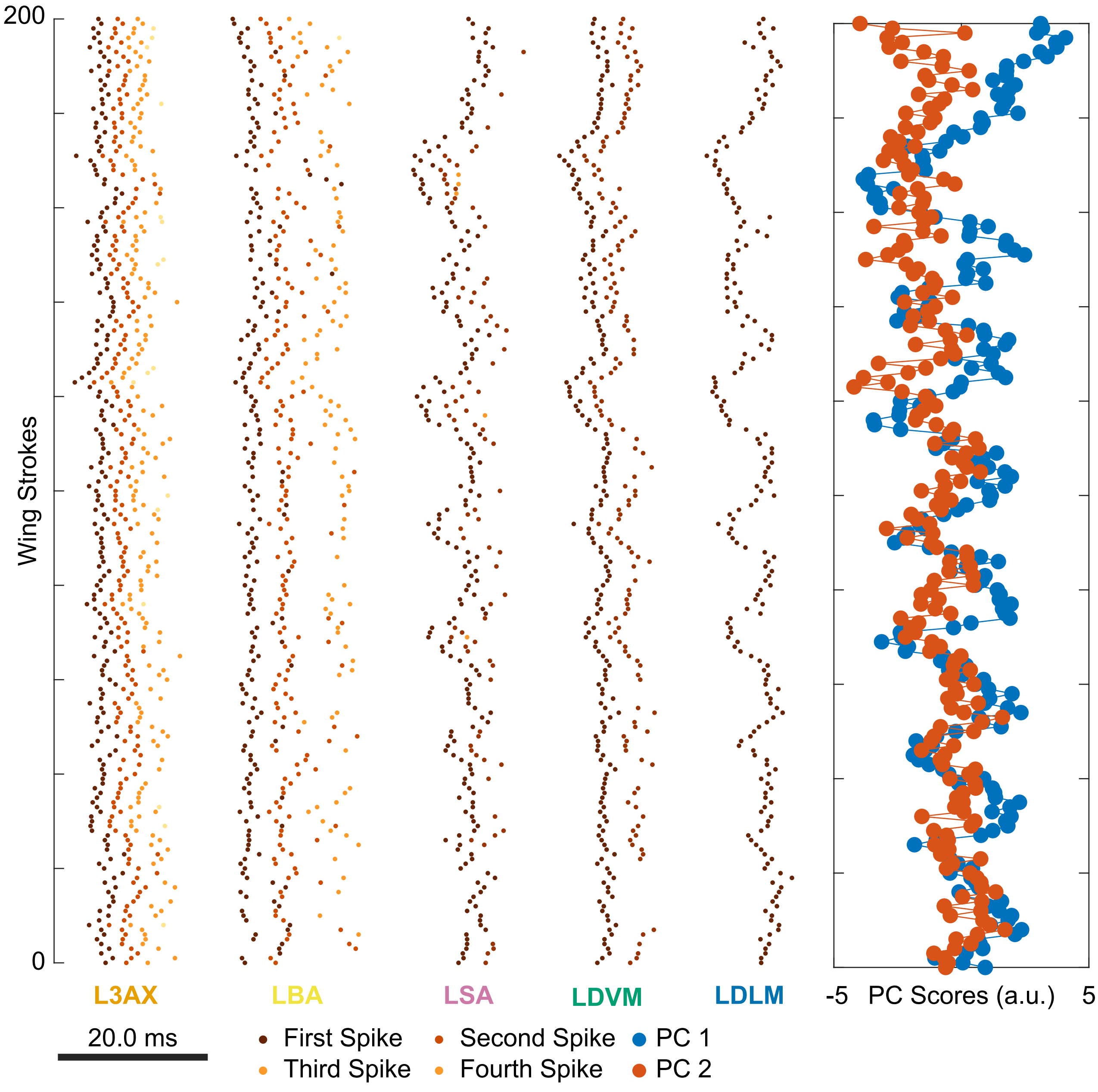

### Supplemental Figure 2

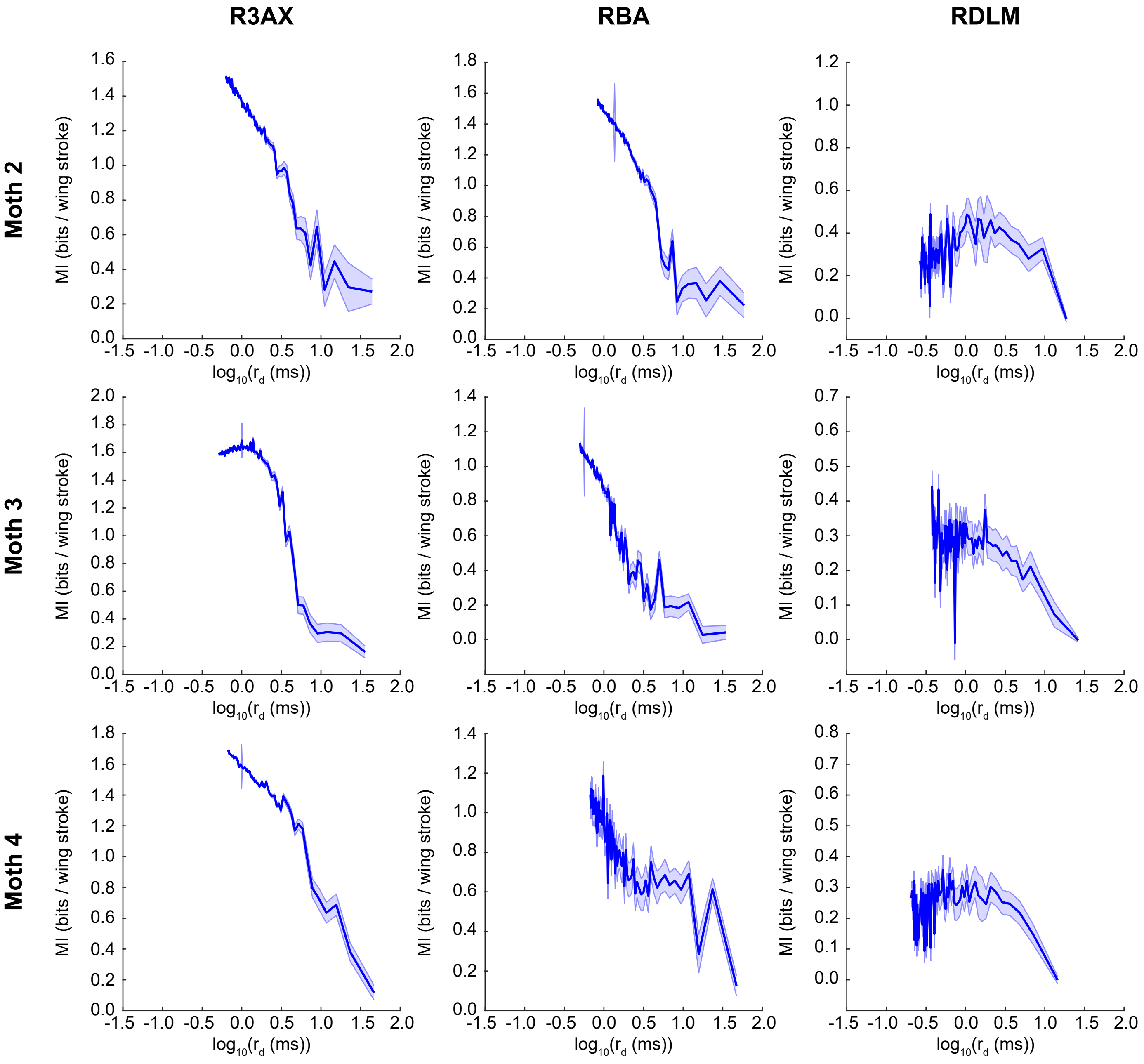

### Supplemental Figure 3

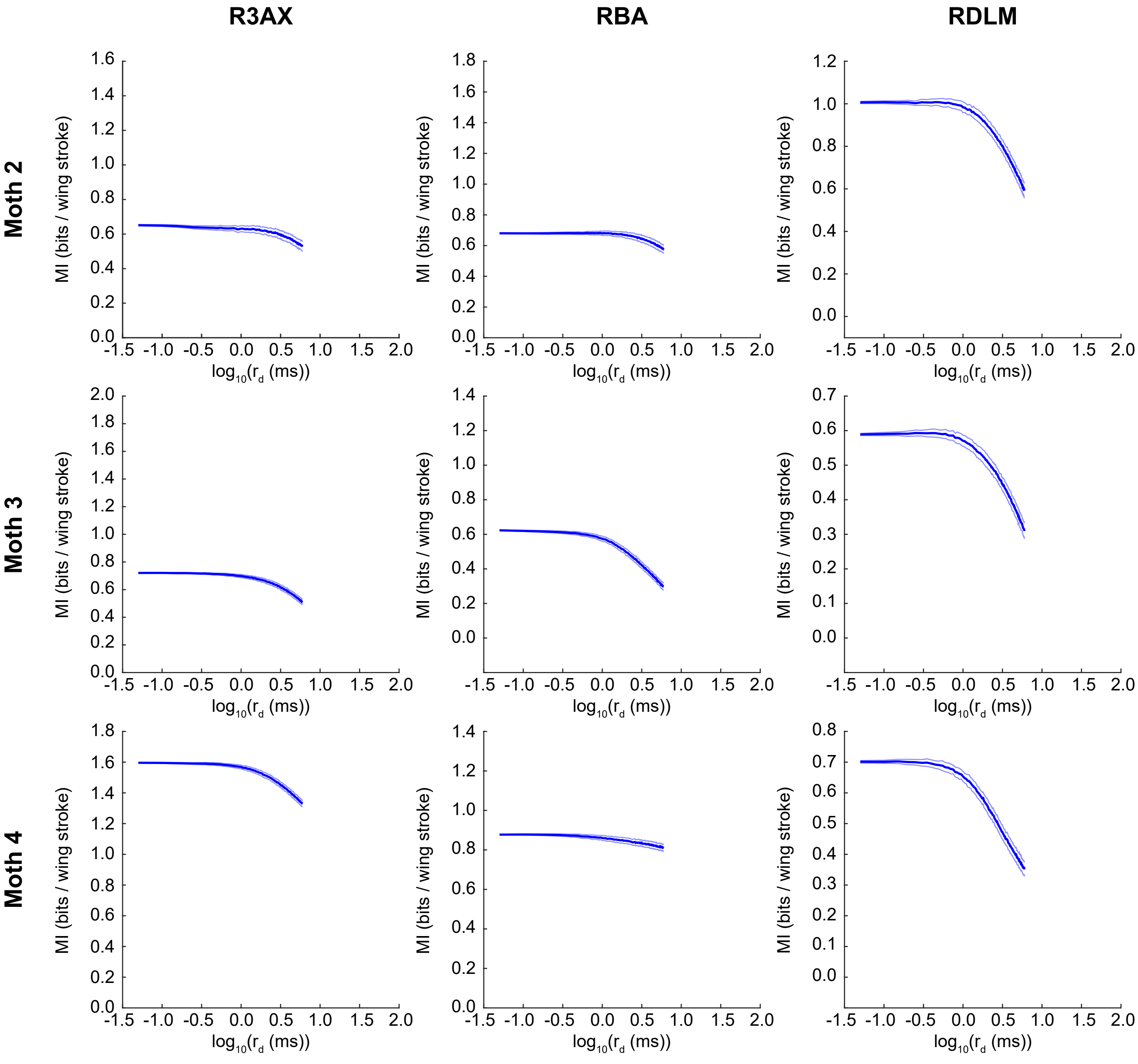
